# Comparative Transcriptomic Analysis of ATRA-Resistant and ATRA-Sensitive APL Cell Lines Identifies lncRNA Biomarkers Associated with Drug Resistance

**DOI:** 10.64898/2026.01.27.702191

**Authors:** Oviya Marimuthu, Shinde Nikhil, Raja Natesan Sella

## Abstract

Acute promyelocytic leukemia is a distinct subtype of acute myeloid leukemia characterized by the t(15;17) translocation, leading to the PML (Promyelocytic leukemia protein)-RARA (Retinoic Acid Receptor Alpha) fusion protein. Although *PML-RARA* fusion is common, there are 20 more fusion events also reported in APL. All -trans retinoic acid (ATRA) is a standard drug for APL, leading to significant improvement in patient outcomes; nevertheless, a small fraction of patients still experience relapse, and some patients exhibit resistance to the drug. Long non-coding RNAs (LncRNAs) are recognized as promising biomarkers for cancer diagnosis, prognosis, and treatment response. In this study, we used ATRA-Resistant (AP1060) and ATRA -Sensitive (NB4), both treated and untreated cell line transcriptomic data retrieved from the NCBI Gene Expression Omnibus(GEO) database to perform transcriptomic analysis with bioinformatic tools. We utilized the LncRAnalyzer pipeline to predict the lncRNAs, followed by differential expression analysis using DESeq2. Weighted Gene Co-expression Network Analysis (WGCNA) was employed to construct lncRNA co-expression modules associated with ATRA resistance. BEDTools is used to identify cis-acting target genes of lncRNAs.LncRNA -miRNA sponging identified by miRanda algorithm. The identified miRNAs reveal their significant role in APL and other leukemia subtypes. The results of the study show that the identified lncRNAs from the miRNA-LncRNA network are promising biomarkers for ATRA resistance.

## INTRODUCTION

Acute promyelocytic leukemia (APL) is a distinct subtype of acute myeloid leukemia (AML) accounts for about 10-15% of AML. A rare type of blood cancer was reported and described in 1957 by Norwegian hematologist, LK Hillested.APL is characterized by reciprocal translocation between the *PML*(Promyelocytic leukemia protein) gene on chromosome 15 and the RARα(Retinoic Acid Receptor Alpha) gene on chromosome 17, denoted as t(15;17)(q22;q21), leading to the formation of fusion genes (Jimenez et al., 2020). *PML-RARA* fusion gene blocks the differentiation of promyelocytes and increases the proliferation of the immature blasts. To treat this, all-trans retinoic acid (ATRA) was introduced as a differentiation agent for promyelocytes and, in turn, it improved the remission rates of APL patients. However, 10 -20% of APL patients develop relapse and resistance to ATRA, limiting its clinical effectiveness (S. et al., 2026; Sun et al., 2004). Therefore, identification of molecular biomarkers that predict the response to ATRA is essential for effective clinical use.

Long non-coding RNAs(lncRNAs) are defined as RNA molecules longer than 200 nucleotides(nt) in length. Functional analysis of lncRNAs demonstrated their diverse structural and regulatory roles in transcriptional and translational levels (Hu et al., 2018). Increasing evidence indicates the role of lncRNAs in leukemia, including differentiation, malignant proliferation, and drug resistance(Yu et al., 2018a). Therefore, understanding the relationship between lncRNAs and drug resistance in malignant tumors helps to develop strategies that can enhance patient outcomes (Bhan et al., 2017).Several studies reported the expression levels of lncRNAs and their correlation with treatment response (Gao et al., 2020). Previous studies have suggested that lncRNAs play a crucial role in myeloid differentiation and APL treatment (Yu et al., 2018b).

In this study, we conducted a thorough transcriptomic analysis utilizing ATRA-resistant (AP1060) and ATRA-sensitive (NB4) APL cell lines under both treated and untreated conditions, employing publicly accessible datasets from the NCBI Gene Expression Omnibus (GEO) database. We employed various bioinformatic tools, such as LncRAnalyzer for predicting lncRNAs and DESeq2 to identify the differentially expressed genes. The cis-acting target genes of lncRNAs were predicted using BEDTools to investigate their genomic regulatory relationships (Augustino et al., 2020) (Shao et al., 2019). Additionally, lncRNA–miRNA interactions were assessed using the miRanda algorithm to establish potential sponging networks. Functional enrichment analysis of the associated target genes revealed the biological pathways involved in ATRA resistance. In summary, this study aims to identify lncRNAs as potential biomarkers in drug resistance.

## Methods

### Datasets Collection and processing

Data Source under different conditions (untreated, TSA, ATRA, TSA + ATRA)

The raw RNA-seq data for **the NB4, AP1060, and HL-60 cell lines** under different experimental groups (+ATRA, +TSA, +ATRA+TSA) and an untreated control were retrieved from the **NCBI Gene Expression Omnibus (GEO)** database (https://www.ncbi.nlm.nih.gov/geo/) under the accession number **GSE175507**, as reported in previous studies (Li et al.,2022). The **human reference genome assembly GRCh38.p14** was downloaded from NCBI datasets for sequence alignment and downstream analyses.

### LncRAnalyzer workflow

To predict the lncRNAs, we used our in-house pipeline called LncRAnalyzer, designed for identifying lncRNAs and novel protein-coding transcripts (NPCTs). The pipeline undergoes a series of computational analyses to predict the lncRNAs. It is incorporated with different tools such as RNAsamba, CPAT, CPC2, LncFinder, and LGC(Nikhil et al., 2025). The final output produces a Venn diagram; we selected the intersection of lncRNAs to enhance the accuracy. We analyzed two datasets, one comprising **NB4 (ATRA-sensitive) cells** with untreated control and ATRA-treated samples, and the other comprising **AP1060 (ATRA-resistant) cells** with untreated control and ATRA-treated samples.

### Differential Expression Analysis

Differential expression analysis of long non-coding RNAs (lncRNAs) was performed using the **DESeq2** package in **R**. DESeq2 performs size factor estimation, dispersion, and the negative binomial distribution as a workflow to test the differential gene expression(Nikhil et al., 2024). In our research, we are focusing on the ATRA drug alone, so we selected NB-4 + Treatment and control, AP1060 + Treatment and control total of 12 samples for further analysis. Raw read count data of lncRNAs were used as input for the analysis. Low-abundance transcripts with read counts below 10 in more than 80% of the samples were filtered out before normalization. Differentially expressed lncRNAs between ATRA-sensitive (NB4) and ATRA-resistant APL samples were identified. Statistical significance was determined based on the Benjamini–Hochberg adjusted p-value (False Discovery Rate, FDR), with an adjusted p-value (FDR) < 0.05 and |log_2_ fold change| ≥ 1 considered as the threshold for significant differential expression(Love et al., 2014).

### WGCNA NETWORK construction and Module identification

Network construction and visualization of samples were performed using the WGCNA package. Gene counts are normalized, and low-expressed genes are removed. The determination of the adjacency matrix for a scale-free topology involved selecting a soft threshold power, which was analyzed by the pickSoftThreshold function in WGCNA (Saris et al., 2009). Then, a hierarchical clustering and modules with a similarity of 0.8 are merged. The correlation of each gene module was analyzed, and the significance of each gene was calculated (Yuan et al., 2025).

In the present study, WGCNA was applied to construct a gene co-expression network using expression data from ATRA-resistant (AP1060) and ATRA-sensitive (NB4) acute promyelocytic leukemia (APL) cell lines. The analysis identified distinct co-expression modules. Among these, the most significant module was selected based on a module–trait correlation analysis using clinical characteristic data, allowing for the identification of key genes and lncRNAs potentially involved in ATRA drug resistance and sensitivity.(Dong et al., 2024)

### LncRNA cis target prediction analysis

To predict the cis target genes of differentially expressed 16 lncRNAs, we used BED-Tool’s window function, which searches for protein-coding genes located within 100kb upstream and downstream of lncRNAs.Genes situated in this region are considered as a potential cis-regulatory target (Liu et al., 2021).(Augustino et al., n.d.). The genomic coordinates of lncRNAs and protein-coding genes were obtained from the **human reference genome GRCh38** (GENCODE annotation).

### Functional Enrichment Analyses of target genes

Biological functions and pathways of differentially expressed lncRNAs target genes in the yellow module were analyzed by Functional enrichment analysis. We used Gene Ontology (GO) and Kyoto Encyclopedia of Genes and Genomes (KEGG) pathway analyses to identify the biological processes, cellular components, and molecular functions of target genes associated with APL (Ke & Ge, 2024).

### LncRNA-miRNA target prediction

Human miRNA sequences (5p/3p specific) were retrieved from miRBase and used to predict interactions with lncRNA transcripts using the miRanda algorithm (Movassagh et al., 2022). miRanda identifies putative miRNA–target interactions based on sequence complementarity and thermodynamic stability. Predictions were performed using a minimum alignment score threshold of 140 and a minimum free energy cutoff of −20 kcal/mol to reduce false positives. Then, we filtered the high-confidence interactions with a Score ≥ 200 and ΔG ≤ −30 kcal/mol, which were kept. For further analysis, interactions were grouped by unique lncRNA identifiers (MSTRG IDs) to identify the miRNAs targeting each lncRNA (Qin et al., 2020).

### miRNA-mRNA target prediction

Human miRNA sequences (5p/3p-specific) were retrieved from miRBase and the human reference mRNA FASTA file derived from the GRCh38 genome assembly. The miRNA-mRNA target prediction was performed using the previously mentioned miRanda–based approach and filtering criteria, as described for the lncRNA-miRNA interaction analysis (Sagot, 2023).

## RESULT

### Data Download, LncRNAs Prediction and Differential Expression Analysis

In this study, we downloaded the expression matrix for the series of **GSE175507** (36 samples) from NCBI. The raw reads are submitted to the pipeline to predict the lncRNAs. The pipeline comprises 7 different tools for predicting lncRNAs. For high accuracy, we selected the lncRNAs predicted by all 7 tools. The LncRAnalyzer tool generated the upset plot as an output; a total of 3186 lncRNAs were predicted by all 7 tools (Fig 1). The quantitative distribution of lncRNAs based on their genomic location is represented in a bar plot (Fig 2) (Chen et al., n.d.).We performed differential gene expression for the 12 samples; as a result, we identified 3693 genes and 208 lncRNAs.

**Figure 1.**
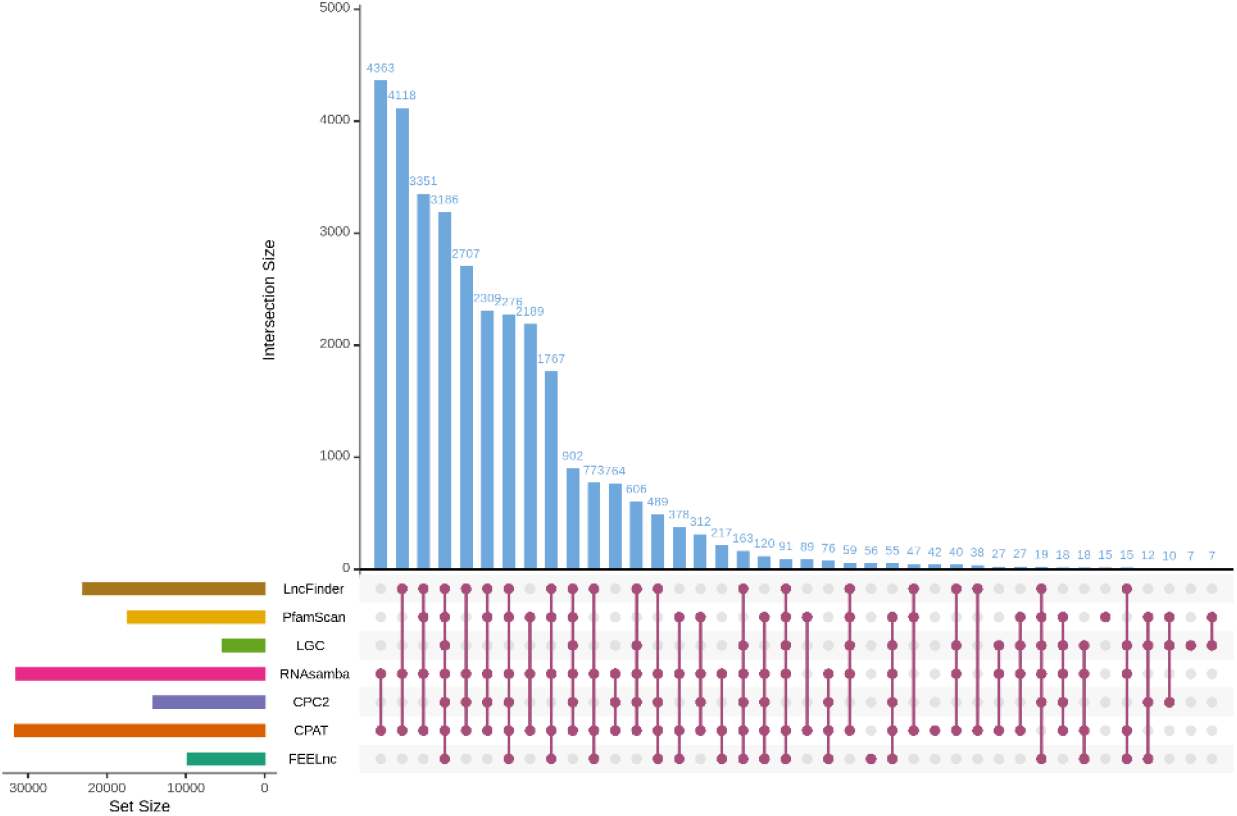
Identified lncRNAs(3186) from the APL cell line dataset by the LncRAnalyzer pipeline

**Figure 2.**
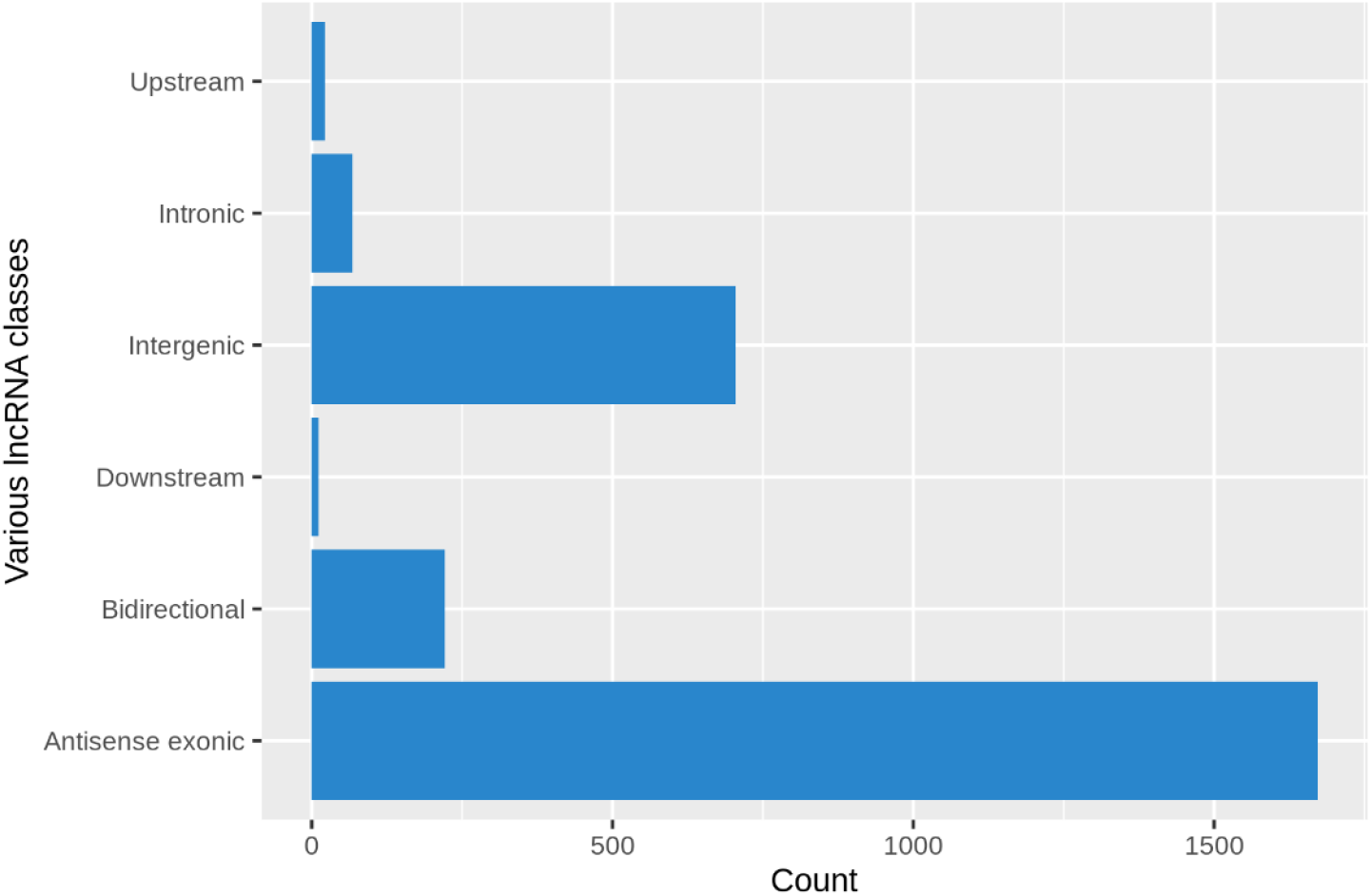
Quantitative distribution of various lncRNA classes identified in the analysis of the APL dataset

### WGCNA Network Construction and Module Identification

WGCNA analysis was performed using the gene expression matrix. Based on gene correlation, the samples were hierarchically clustered. To ensure the scale-free topology, a soft thresholding power was selected, achieving a scale-free fit index (R2) of 0.8, and the soft threshold power was determined to be 28 (Saris et al., 2009) (Fig 3). Based on the power, a gene dendrogram was generated with merged and unmerged modules. The merged module is considered a stable and biologically meaningful module. The merged modules included black, blue, green, grey, red, turquoise, and yellow (Fig 4). To visualize their expression pattern heatmap was plotted(Fig 5). Among these modules, the yellow module showed a significant expression difference between control and treated cell lines. This module consists of 446 genes and 43 lncRNAs. A heatmap was generated for these 43 lncRNAs to examine their expression pattern(Fig 6). To improve the consistency across replicates, the mean expression value of triplicate samples was calculated for each condition. Based on low mean connectivity, we selected 16 Differentially expressed lncRNAs and plotted the heatmap(Fig 7). These lncRNAs are significantly upregulated in AP1060 upon ATRA treatment and downregulated in NB4 upon ATRA treatment.

**Figure 3.**
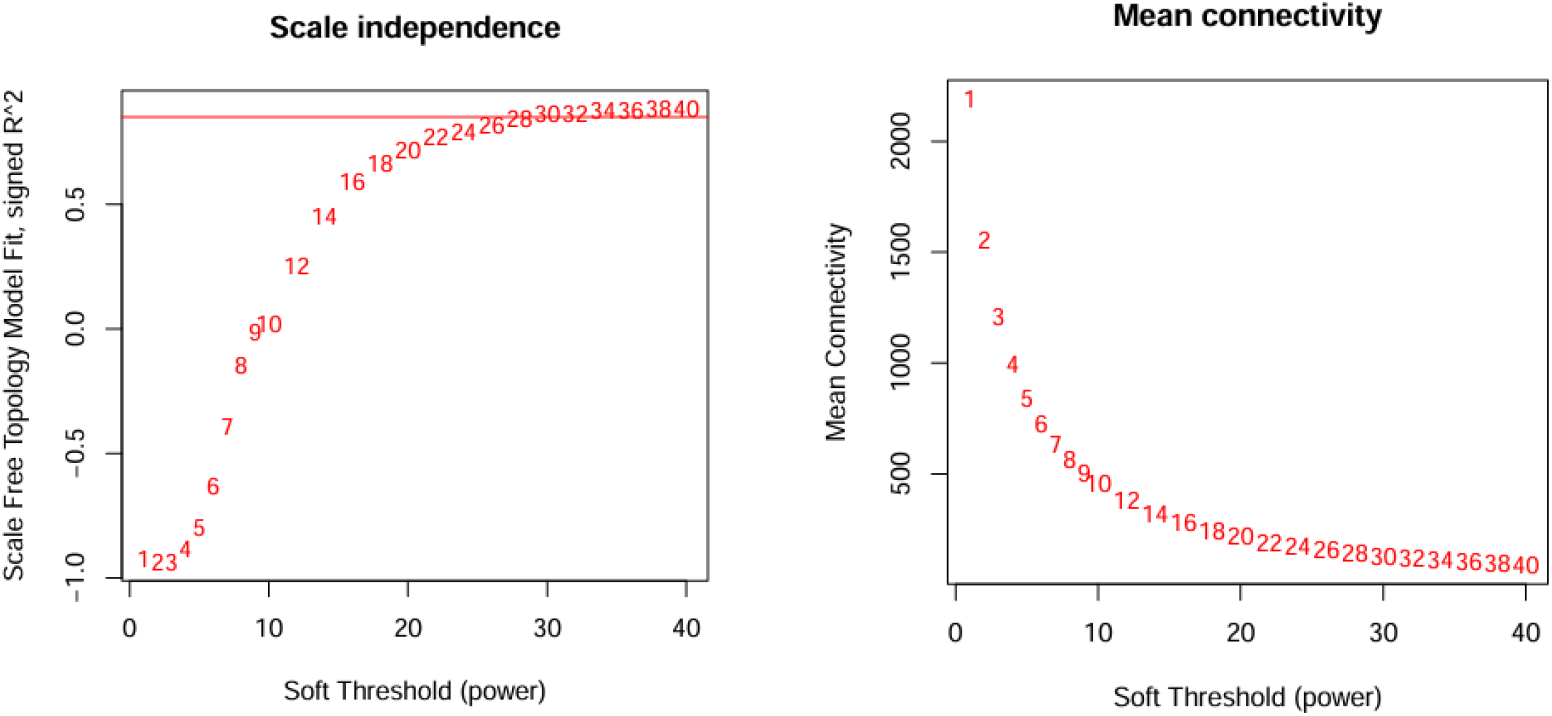
Network topology for 0.8 soft thresholding powers. The mean connectivity value of 28 is selected for plotting the cluster dendrogram.

**Figure 4.**
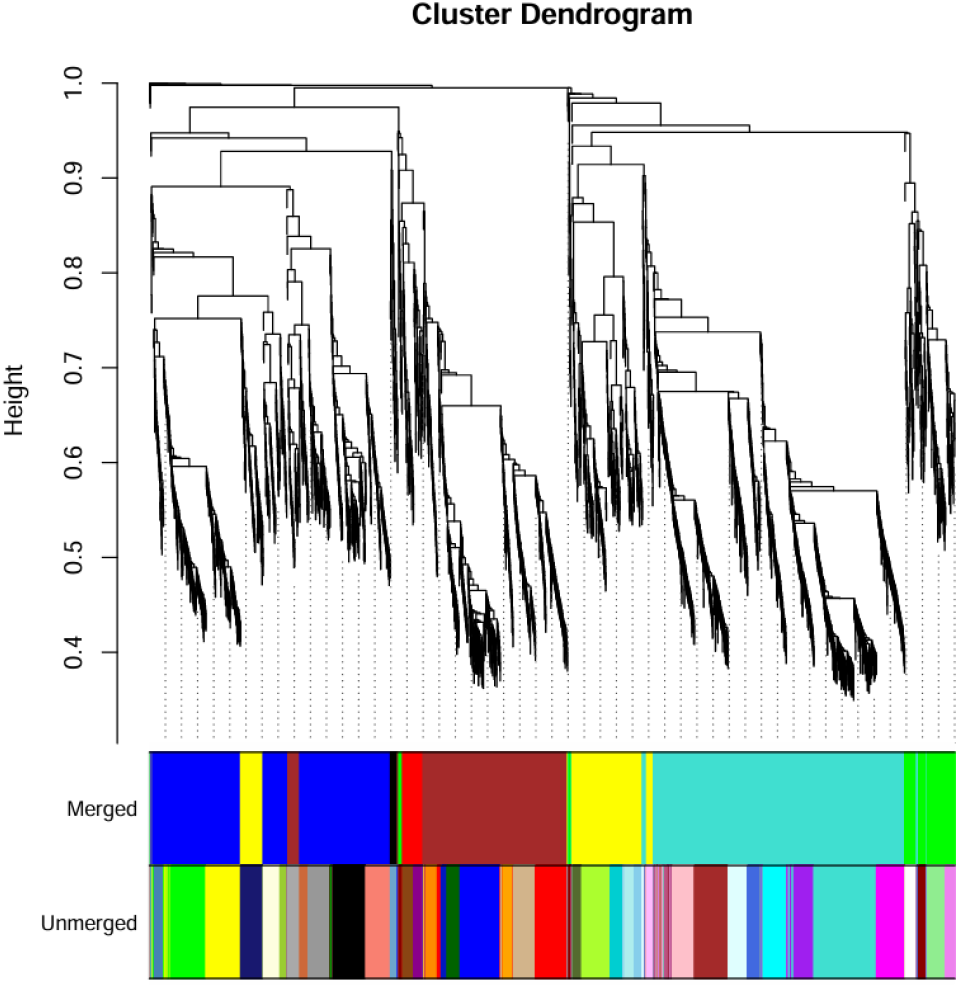
The cluster dendrogram and color display of co-expression network modules for all genes. The short vertical line corresponded to a gene, and the branches corresponded to the co-expressed genes.

**Figure 5.**
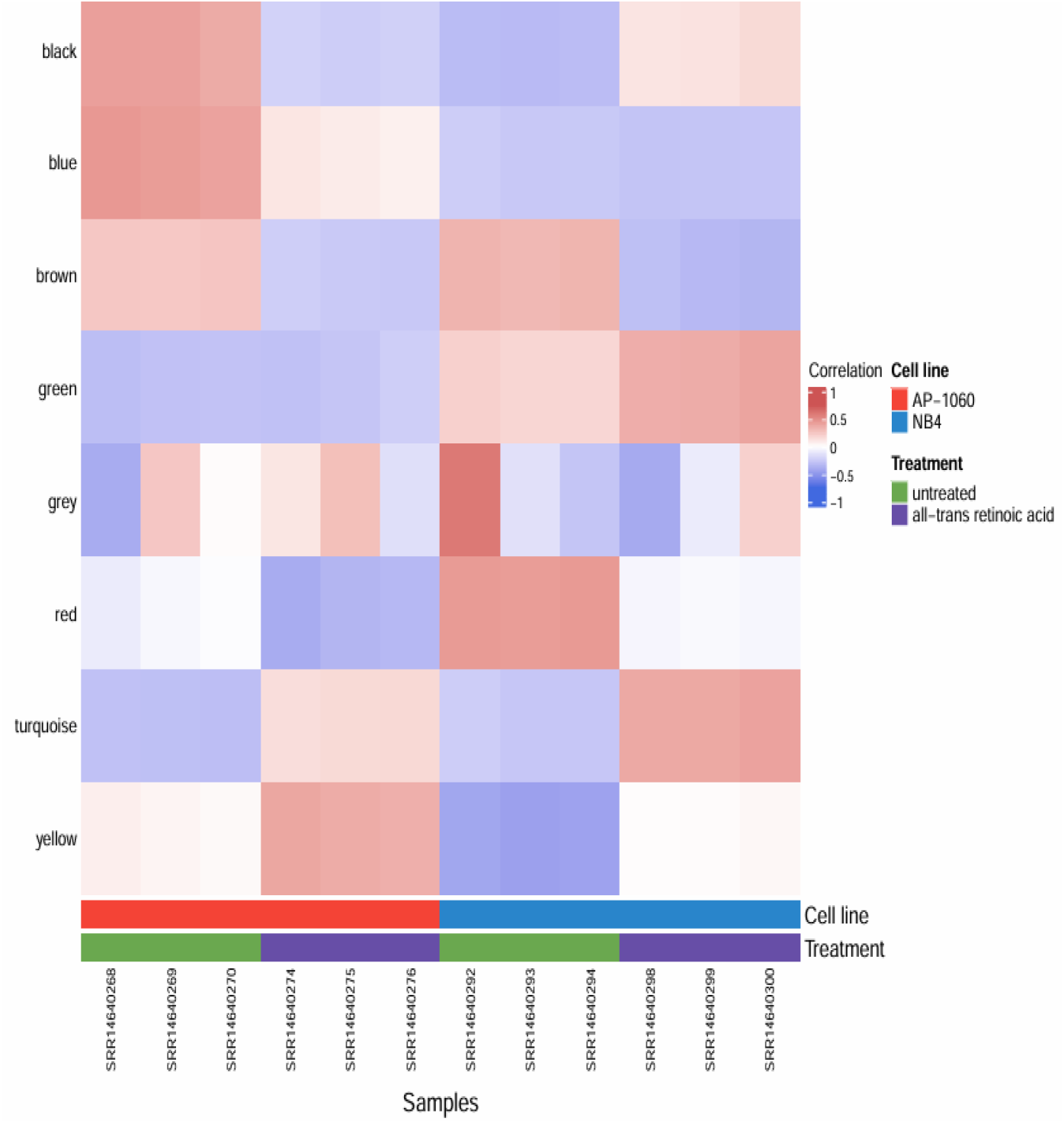
Correlation matrix of module eigengene values obtained from WGCNA

**Figure 6.**
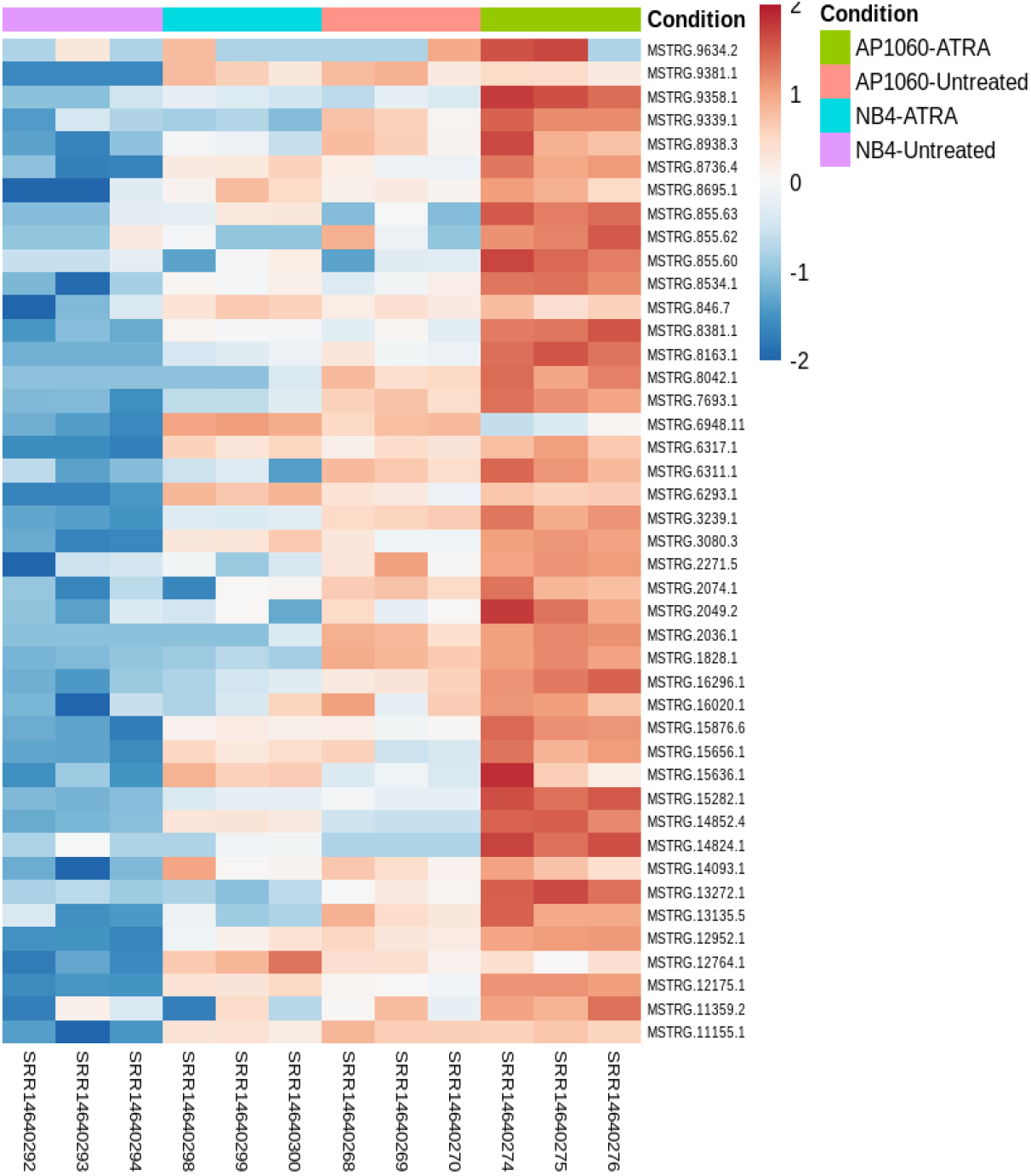
Heatmap showing the expression profiles of 43 differentially expressed lncRNAs across NB4 and AP1060 cell lines under ATRA-treated and untreated conditions. The lncRNAs show higher expression in AP1060 with ATRA treatment and reduced expression in NB4 with ATRA treatment.

**Figure 7.**
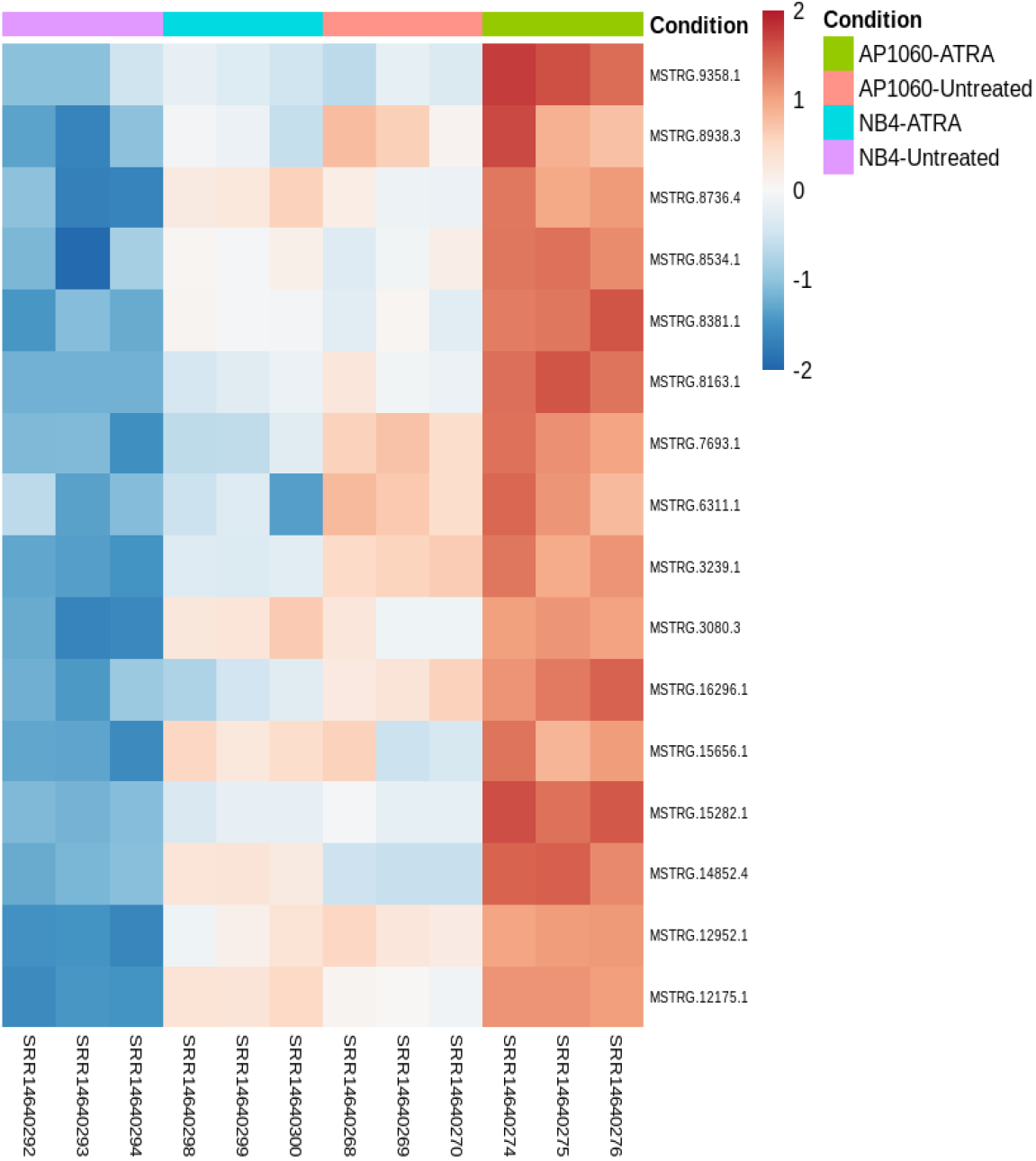
Heatmap illustrating the expression of 16 lncRNAs across NB4 and AP1060 cells under treated and untreated conditions. These lncRNAs are significantly upregulated in AP1060 upon ATRA treatment and downregulated in NB4 upon ATRA treatment. Colors indicate normalized expression values.

### LncRNA cis target prediction

To understand the possible cis-regulatory roles of the differentially expressed lncRNAs, we predicted their nearby target genes using BEDTools (Augustino et al., n.d.). A total of 16 differentially expressed lncRNAs were analyzed to identify protein-coding genes located within 100 kb upstream and downstream of each lncRNA (Liu et al., 2021).Using this approach, we identified 32 protein-coding genes as putative cis-target genes of the 16 lncRNAs. The cis-lncRNA–gene interaction network revealed that 9 lncRNAs were associated with more than one neighboring gene, suggesting their possible involvement in local transcriptional regulation, and 7 lncRNAs showed a one-to-one relationship with protein-coding genes, indicating specific cis-regulatory associations (Fig 9).

**Figure 8.**
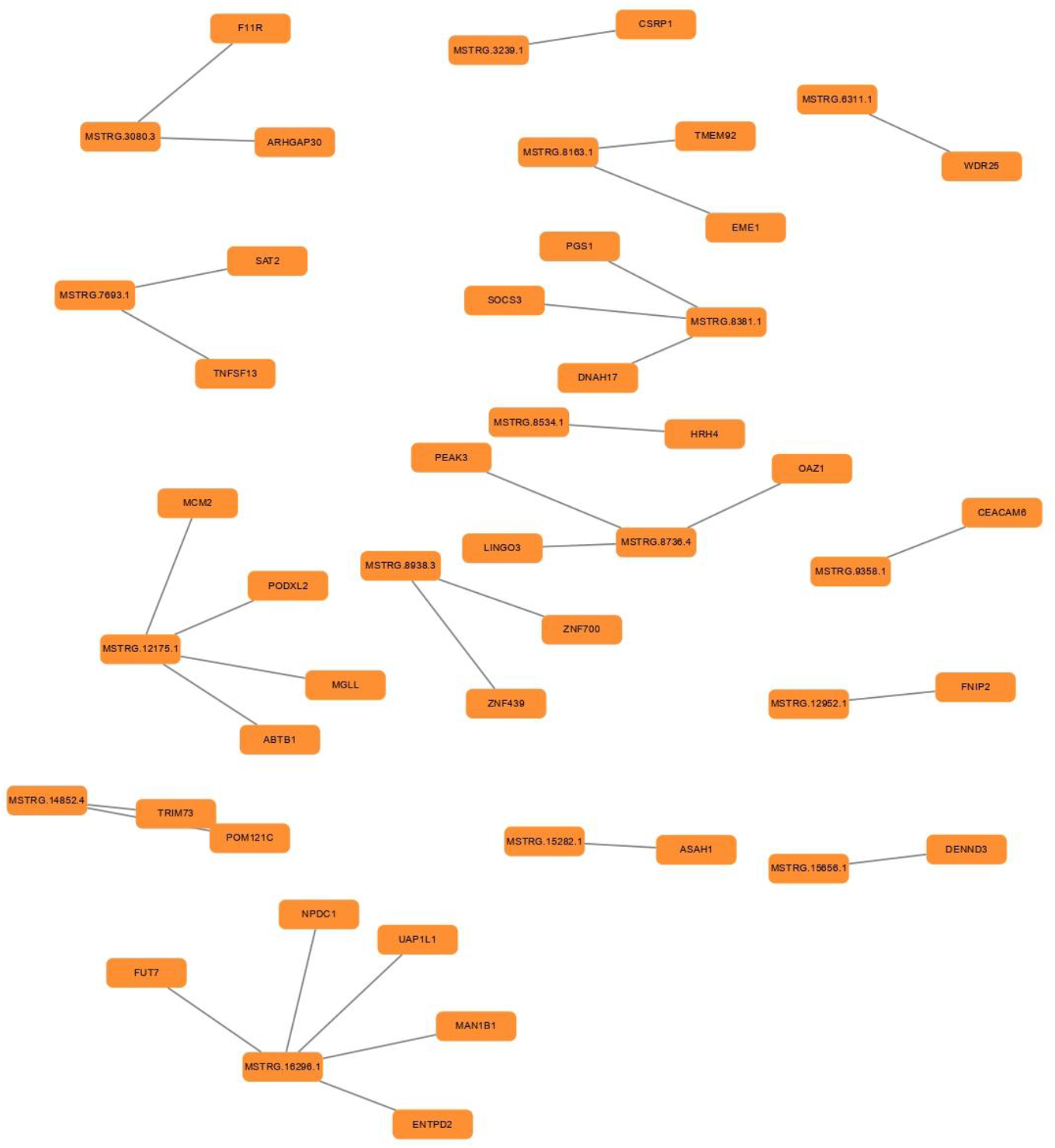
LncRNA–gene interaction network showing the predicted associations between differentially expressed lncRNAs and their corresponding target genes. Each node represents an lncRNA (MSTRG ID), and edges indicate predicted regulatory interactions.

**Figure 9.**
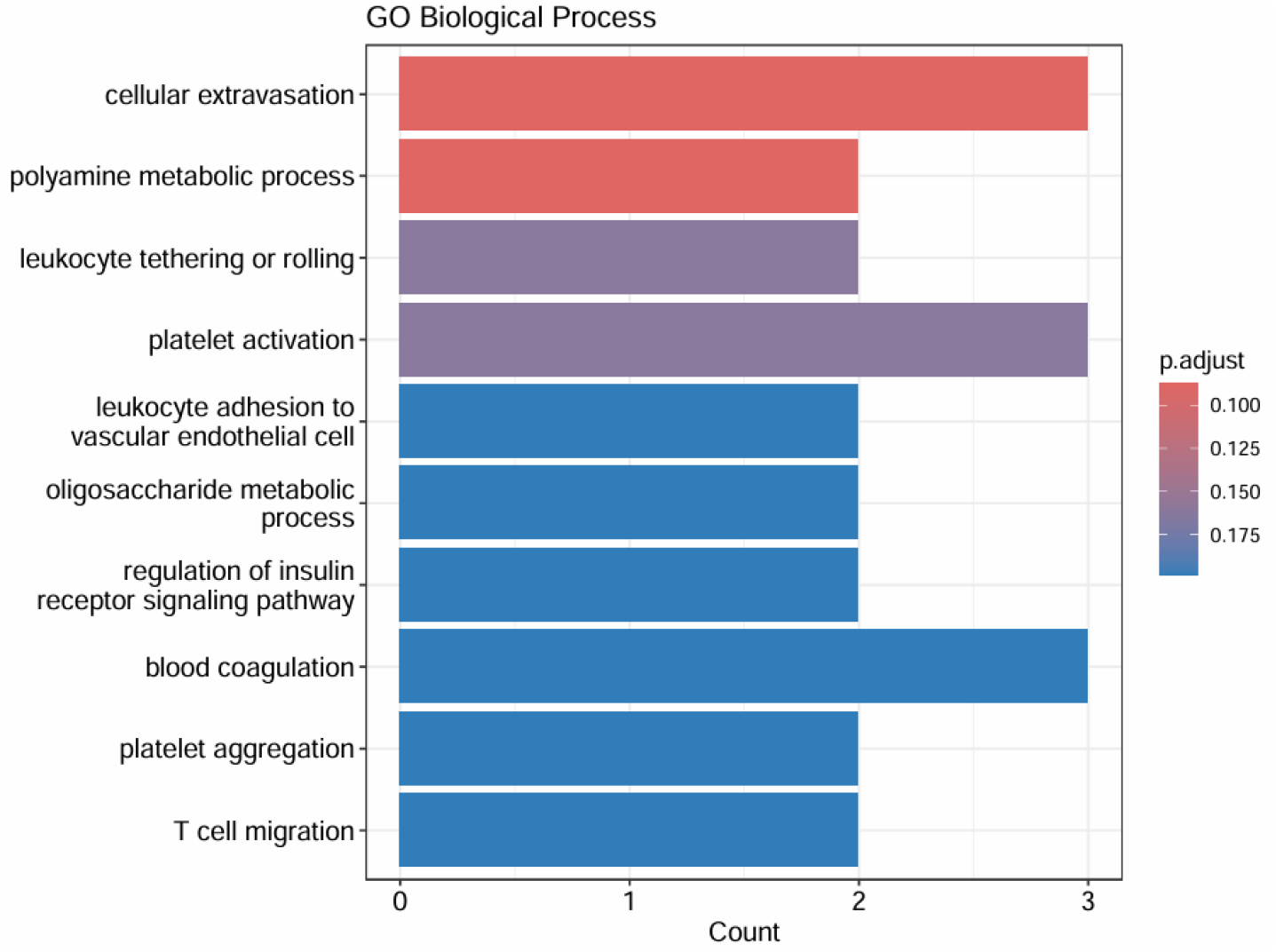
GO Biological process for target genes

### Functional enrichment analysis of target genes

Gene ontology enrichment analysis was performed for the predicted 32 cis-target genes to understand their biological roles(Fig 10). The result showed that these genes were mainly enriched in biological processes related to cell movement, metabolism, and blood -related functions (Ke & Ge, 2024). The significant enrichment was observed in leukocyte tethering and T cell migration. In addition, platelet activation and blood coagulation were also enriched. Together, these results indicate that the cis-target genes of differentially expressed lncRNAs are associated with immune regulation, metabolism, and blood formation.

**Figure 10.**
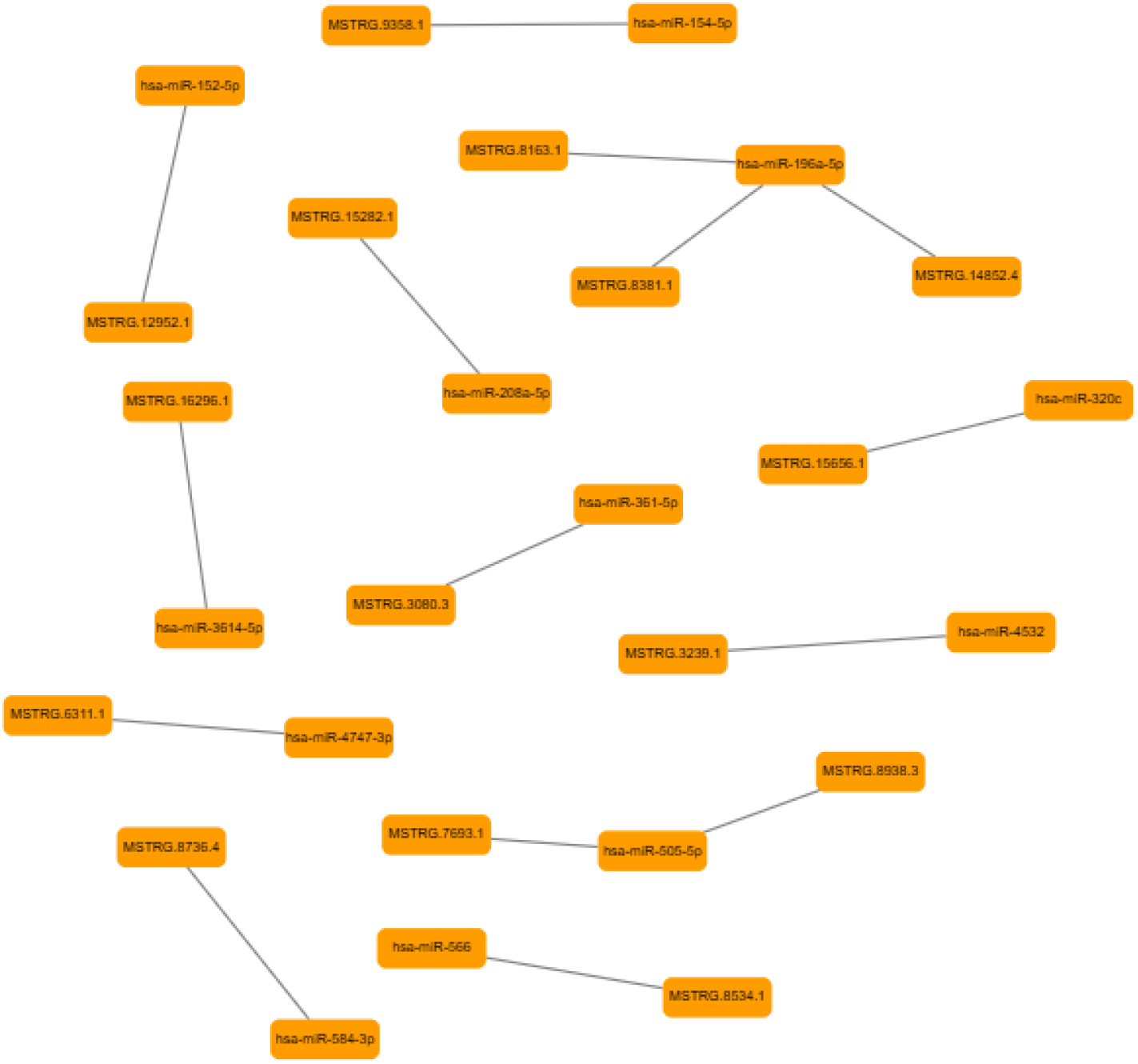
This figure shows the predicted interactions between miRNAs and lncRNAs identified using the miRanda tool. Nodes labeled with hsa**-** represent miRNAs, while nodes labeled with MSTRG represent lncRNA transcripts. Lines connecting the nodes indicate predicted miRNA– lncRNA binding interactions based on sequence complementarity. The network highlights potential regulatory relationships between miRNAs and lncRNAs.

### LncRNA-miRNA target prediction

LncRNA -miRNA targets were identified by the miRanda tool. Based on the mentioned criteria, we identified 63 high-confidence interactions, involving 19 unique miRNAs and 16 unique lncRNAs (MSTRG transcripts). And we constructed the network, which reveals key regulatory hubs within the interaction network(Fig 11). The most targeted transcripts are MSTRG.8736.4 and MSTRG.14852.4, each interacting with 10 different miRNAs. In addition, hsa-miR-505-5p and hsa-miR-196a-5p were identified as major miRNA regulators, each targeting multiple transcripts.

### miRNA-mRNA target prediction

miRNA-mRNA targets were identified by the miRanda tool (Riolo et al., 2021). miRNAs that were previously identified as targeting lncRNAs were subsequently selected, and their corresponding mRNA targets were examined. By integrating miRNA, lncRNA, and mRNA interaction data, we mapped miRNA–lncRNA–mRNA regulatory relationships and identified 13 unique putative interactions (Table 1). The miRNA was predicted to target lncRNA and gene, suggesting that these lncRNAs may modulate gene expression by sequestering miRNA.These interactions highlight the potential role of lncRNAs in post-transcriptional gene regulation by acting as miRNA sponges.

**Table 1.** The table summarizes the predicted regulatory relationships among microRNAs (miRNAs), long non-coding RNAs (lncRNAs), and their putative target protein-coding genes.

| miRNA | lncRNA | Gene |
| --- | --- | --- |
| hsa-miR-1251-3p | MSTRG.12175.1 | *RPS6KA1* |
| hsa-miR-152-5p | MSTRG.12952.1 | *IBA57* |
| hsa-miR-154-5p | MSTRG.9358.1 | *MTUS2* |
| hsa-miR-196a-5p | MSTRG.14852.4 | *HOXB3* |
| hsa-miR-196a-5p | MSTRG.8163.1 | *HOXB3* |
| hsa-miR-196a-5p | MSTRG.8381.1 | *HOXB3* |
| hsa-miR-208a-5p | MSTRG.15282.1 | *N4BP3* |
| hsa-miR-320c | MSTRG.15656.1 | *ACAN* |
| hsa-miR-3614-5p | MSTRG.16296.1 | *WDR81* |
| hsa-miR-361-5p | MSTRG.3080.3 | *LILRA6* |
| hsa-miR-4532 | MSTRG.3239.1 | *C19orf85* |
| hsa-miR-4747-3p | MSTRG.6311.1 | *KCNAB2* |
| hsa-miR-505-5p | MSTRG.7693.1 | *DCAF5* |
| hsa-miR-505-5p | MSTRG.8938.3 | *DCAF5* |
| hsa-miR-566 | MSTRG.8534.1 | *SH3BGRL2* |
| hsa-miR-584-3p | MSTRG.8736.4 | *MOV10* |

## Discussion

Acute promyelocytic leukemia (APL) is a rare and severe type of blood cancer that constitutes approximately 10–12% of acute myeloid leukemia cases (Li et al., 2022). All-trans retinoic acid (ATRA) is a standard drug administered to individuals diagnosed with APL.ATRA treatment has significantly enhanced the patient survival rate (Liang et al., 2021) (Iyer et al., 2023). Nonetheless, a group of patients experience relapse and develop resistance to ATRA, which continues to pose a significant challenge in APL treatment (Jimenez et al., 2020). Identifying molecular biomarkers to predict treatment resistance is crucial for improving patient outcomes. The present study analyzed the novel differentially expressed lncRNAs between ATRA-resistant (AP1060) and ATRA-sensitive (NB4) cell lines (Sun et al., 2004). The DESEq2 package was used to identify the differentially expressed lncRNAs. As a result, we identified 3693 genes and 208 lncRNAs.And we performed WGCNA analysis to identify co-expressed gene modules; we identified 7 modules: black, blue, green, grey, red, turquoise, and yellow modules. Among these modules, the yellow module showed a significant expression difference between control and treated cell lines (Love et al., 2014). We plotted a heatmap to visualize the expression pattern. We selected the top 16 differentially expressed novel lncRNAs based on low mean connectivity between triplicate. And we identified the target genes for those lncRNAs by BEDTool (Liu et al., 2021).Gene ontology reveals the target genes’ role in several biological pathways, such as platelet activation, blood coagulation, and T cell migration, which are closely related to APL (Dong et al., 2024). These findings suggest that our lncRNAs may play a significant role in APL. To further explore the lncRNA, we performed lncRNA-miRNA interaction using the miRanda tool; the output produced 16 high-confidence interactions, involving 16 unique miRNAs and 16 unique lncRNAs (MSTRG transcripts). And we constructed the network for finding regulatory hubs within the interaction network. Followed by that, we looked for miRNA-mRNA targets using the miRanda tool (Fan et al., 2025). miRNAs that were previously identified as targeting lncRNAs were subsequently selected, and their corresponding mRNA targets were examined. By integrating miRNA, lncRNA, and mRNA interaction data, we mapped miRNA–lncRNA–mRNA regulatory relationships and identified 16 unique putative interactions. The miRNA was predicted to target lncRNA and gene, and these lncRNAs may modulate gene expression by sequestering miRNA. These interactions highlight the potential role of lncRNAs in post-transcriptional gene regulation by acting as miRNA sponges.

## Conclusion

The present study provides a comparative transcriptomic analysis of ATRA-resistant (AP1060) and ATRA-sensitive (NB4) APL cell lines. We identified 16 novel lncRNAs exhibiting a differential expression pattern in response to ATRA treatment by integrating differential expression analysis, co-expression network modeling, cis-target prediction, and miRNA-lncRNA-mRNA interaction analyses. The functional enrichment of their target genes suggests a role in hematopoiesis and cellular migration, which are closely linked to APL etiology and treatment efficacy. The results indicate that these lncRNAs may serve as a potential biomarker for ATRA resistance for future experimental validation and therapeutic exploration in APL.

